# Balance between DNA repair, LINE1 suppression and lifespan in mice with SIRT6 Serine 10 phosphorylation site mutations

**DOI:** 10.64898/2026.02.06.704345

**Authors:** Zhihui Zhang, Matthew Simon, Yuan Liang, Alexander Tyshkovskiy, Mila Kaplan, Victoria Paige, Seyed Ali Biashad, Vadim Gladyshev, Andrei Seluanov, Vera Gorbunova

## Abstract

Sirtuin 6 (SIRT6) is an important regulator of DNA repair, metabolism, chromatin maintenance and longevity. SIRT6 Serine 10 phosphorylation controls SIRT6 recruitment to the sites of DNA damage. To explore the effect of SIRT6 Serine 10 phosphorylation on lifespan, we generated two SIRT6 mutant mouse strains: phospho-null S10A and phosphomimetic S10E. The S10E mutant mice demonstrated enhanced DNA repair capacity, elevated LINE1 expression and reduced lifespan in male mice compared to the wild-type and S10A mice. This result suggests that SIRT6 S10E mutation enhances DNA repair capacity at the expense of reduced LINE1 silencing leading to shorter lifespan. While both SIRT6 functions in DNA repair and chromatin maintenance are important for longevity, our results suggest that when the balance between these functions is shifted, diminished of LINE1 control has a stronger impact on lifespan than enhanced DNA repair.

## Introduction

Sirtuins, a highly conserved family of NAD+-dependent deacetylases, play a pivotal role in regulating diverse biological pathways. Initial findings that Sir2 regulates yeast lifespan have spurred significant interest in mammalian sirtuins as potential targets for longevity research(Guarente, 2000; Lin *et al*, 2000). Among the seven sirtuin genes, SIRT6 has emerged as a key regulator of longevity. Multiple studies have generated SIRT6 knockout (KO) mice, reporting reduced lifespans, although the severity of phenotypes varies depending on genetic background(Mostoslavsky *et al*, 2006; Simon *et al*, 2019; Xiao *et al*, 2010). While gene knockouts often lead to various pathologies and shortened lifespans, SIRT6 is unique in that its overexpression has been shown to extend lifespan(Kanfi *et al*, 2012). In a study by Roichman et al., SIRT6 overexpression under a CAG promoter resulted in a 27% increase in median lifespan for male mice and 15% for female mice compared to wild-type littermates. Additionally, SIRT6 overexpression enhanced organismal homeostasis and improved multiple aspects of healthspan(Roichman *et al*, 2021).

SIRT6 plays a crucial role in DNA repair, facilitating the repair of both single-strand breaks (SSBs) and double-strand breaks (DSBs)(Mao *et al*, 2011). Mice lacking SIRT6 (Sirt6 KO) exhibit chromosomal abnormalities consistent with defects in DNA repair(Han *et al*, 2015; Mostoslavsky *et al*., 2006). Age-associated heterochromatin reorganization leads to the derepressing of transposable elements, genomic elements that comprise nearly half of the human genome and contribute to genomic instability during aging(De Cecco *et al*, 2019b). Of particular relevance are Type 1 long interspersed nuclear elements (LINE1s), whose retrotransposition activity increases in aged somatic tissues. Activation of LINE1 elements results in the accumulation of cytoplasmic LINE1 DNA, which is detected by the DNA sensor cyclic GMP-AMP (cGAMP) synthase (cGAS)(De Cecco *et al*., 2019b; Simon *et al*., 2019). This detection triggers an innate immune response via activation of the stimulator of interferon genes (STING) pathway, leading to type I interferon (IFN) production and promoting sterile inflammation. Sirt6 KO mice exhibit a significant increase in LINE1 cDNA, reaching levels comparable to or exceeding those seen in aged wild-type mice. In contrast, SIRT6 overexpression suppresses LINE1 activity(Simon *et al*., 2019).

SIRT6 is a highly post-translationally modified protein, with its functions regulated by these modifications. Although the regulation of SIRT6 by those modifications is not extensively studied, our previous research demonstrated that SIRT6 is phosphorylated by JNK at Serine 10 (S10) in response to oxidative stress. This phosphorylation at S10 is essential for the efficient recruitment of SIRT6 to sites of DNA damage(Van Meter *et al*, 2016). To assess the role of SIRT6 S10 phosphorylation on mouse lifespan, we employed CRISPR/Cas9 to generate SIRT6 S10 mutant strains. The S10E mutant exhibited enhanced DNA repair capacity and greater resistance to whole-body gamma irradiation. However, despite enhanced DNA repair, the SIRT6 S10E mutant did not extend lifespan of the mice possibly due to reduced capacity of suppressing LINE1 transposon expression.

## Results

### SIRT6 S10E mutant mice are resistant to DNA damage

Our previous study demonstrated that phosphorylation of SIRT6 Serine 10 residue regulates the DNA repair capacity of fibroblasts by promoting SIRT6 recruitment to DNA double strand break sites(Van Meter *et al*., 2016). To investigate the *in vivo* role of SIRT6 Serine 10 phosphorylation, we utilized the CRISPR/Cas9 system to generate two SIRT6 mutants in mice: a phospho-null Serine-to-Alanine (S10A) mutation and a phosphomimetic Serine-to-Glutamate (S10E) mutation (**Fig. 1A**).

**Figure 1.**
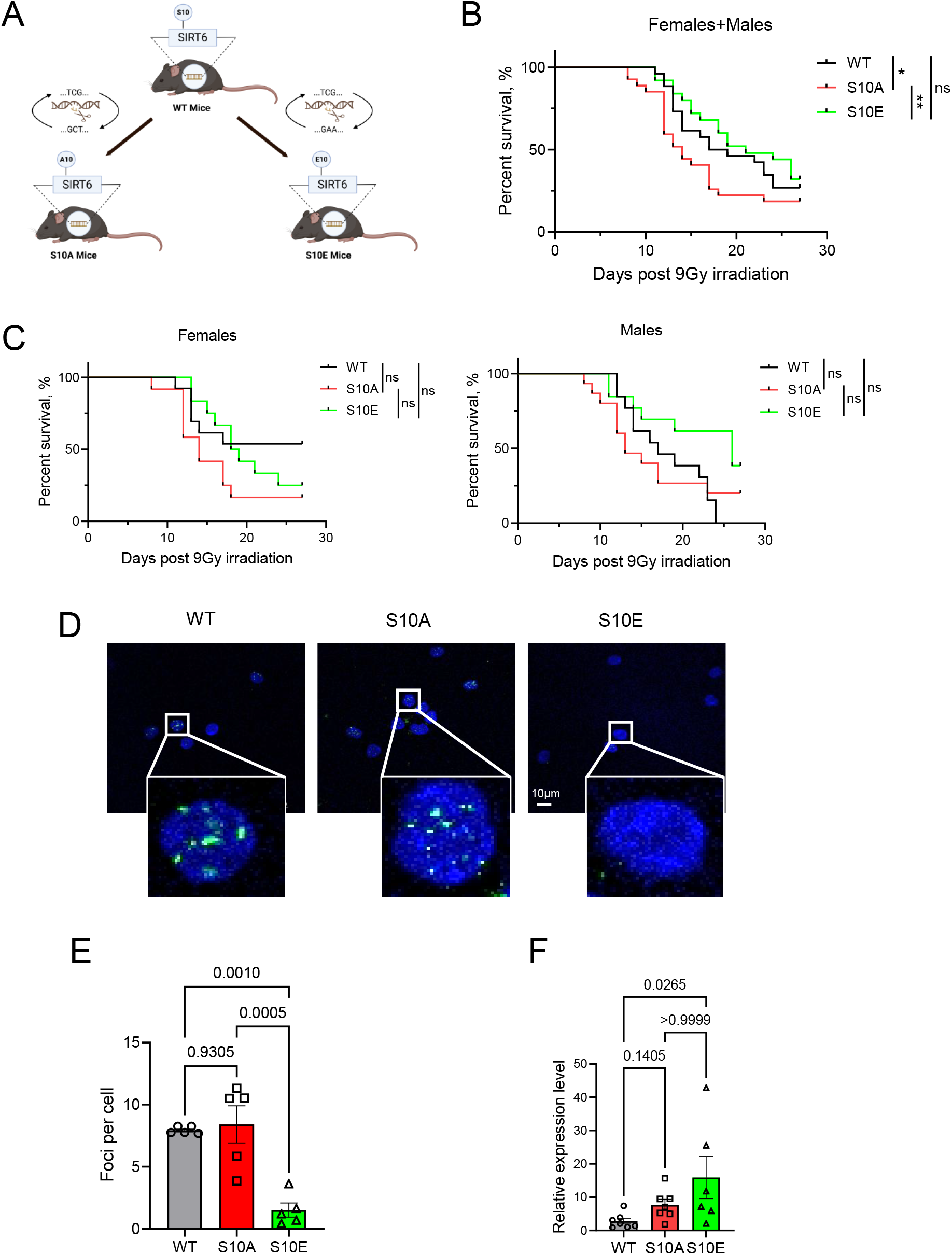
Resistance to irradiation and DNA repair capacity in SIRT6 S10A and S10E mice. **A.** Schematic representation of the CRISPR–Cas9 strategy used to generate Sirt6 S10A (phospho-null) and S10E (phosphomimetic) knock-in mice on the C57BL/6J background. The Ser10 residue in the N-terminus of SIRT6 was replaced by alanine (A10) or glutamate (E10), respectively, to impair or mimic phosphorylation. **B** Survival curves of wild-type (WT), S10A and S10E mice exposed to a single sublethal dose of 9 Gy whole-body γ-irradiation at young adult age (n = 25-27 mice per genotype; sexes combined). S10A mice show reduced post-irradiation survival compared with WT, whereas S10E mice display a trend toward improved survival. Survival differences were assessed using the Gehan–Breslow–Wilcoxon test. *:P<0.05, **P<0.01. **C.**Sex-stratified survival curves for female ((n = 11-13 mice per genotype, left) and male ((n = 12-14 mice per genotype, right) mice after 9 Gy γ-irradiation. The protective effect of the S10E allele is most pronounced in males, whereas S10A males show reduced resistance compared with WT. Survival statistics were calculated as in **D.** Representative confocal images of γH2AX immunofluorescence in peripheral blood mononuclear cells (PBMCs) isolated 72 h after irradiation from WT, S10A and S10E mice. Nuclei are counterstained with DAPI (blue); γH2AX foci marking DNA double-strand breaks are shown in green. Insets show higher-magnification views of individual nuclei. Scale bar, 10 μm. **E.** Quantification of γH2AX foci per nucleus in PBMCs 72 h after irradiation (n = 5 mice per genotype, 50– 70 nuclei scored per mouse). Each symbol represents one biologically independent mouse; horizontal lines indicate mean values and error bars denote s.e.m. S10E PBMCs show significantly fewer γH2AX foci than WT and S10A PBMCs, consistent with enhanced DNA repair, whereas S10A PBMCs retain more residual damage. P values were calculated using one-way ANOVA with post-hoc multiple comparisons and are indicated in the graph. **F.** Relative LINE1 (L1) transcript abundance in PBMCs 72 h after irradiation, measured by RT–qPCR and normalized to Actb (n = 6-7 mice per genotype). Each symbol represents one mouse; bars show mean ± s.e.m. S10E mice display elevated L1 expression compared with WT and S10A mice, indicating that the phosphomimetic mutation compromises SIRT6-mediated LINE1 repression under genotoxic stress. P values were determined by one-way ANOVA; exact P values are shown in the panel.

To assess whether phosphorylation of this residue influences DNA repair capacity *in vivo*, mutant and WT control C57BL/6 mice were exposed to a sublethal dose of 9 Gy γ-irradiation. This dose resulted in 75% lethality across all three genotypes. In agreement with our previous *in vitro* findings, S10A mutant mice exhibited reduced resistance to irradiation and lower survival rates compared to S10E and WT mice. S10E mice exhibited a trend of improved survival relative to wild-type mice (**Fig. 1B**), particularly the male mice (**Fig. 1C**).

To assess DNA repair efficiency in S10E mutants, we isolated peripheral blood mononuclear cells (PBMCs) from irradiated mice and performed γH2AX staining (**Fig. 1D**). Consistent with the survival trends, S10E mice displayed enhanced DNA repair capacity, as indicated by significantly fewer γH2AX foci in PBMCs compared to S10A and WT mice, which showed more residual foci (**Fig. 1D, E**).

### SIRT6 S10E mice show elevated expression of LINE1 elements

In our previous work, we found that unphosphorylated SIRT6 at the S10 residue localizes to the promoters of the retrotransposable element LINE1 (L1) and represses its expression under stress(Van Meter *et al*, 2014) . To evaluate LINE1 expression in S10 mutant mice we performed qRT-PCR on PMBCs from control and irradiated mice. Consistent with the role of S10 phosphorylation in recruiting SIRT6 to DSB sites and away from LINE1 transposons, irradiated S10E mice exhibited significantly higher LINE-1 expression compared to the WT mice. PBMCs from S10A mutants also showed a trend towards increased LINE-1 mRNA levels, possibly due to S10A mutation partially interfering with SIRT6 function, although this increase was not statistically significant compared to both S10E and Bl6 mice (**Fig. 1F**). In summary, the S10E mutant mice demonstrated enhanced DNA repair capacity but reduced suppression of LINE1 expression following irradiation.

We showed previously that elevated LINE1 expression leads to the formation of cytoplasmic LINE1 DNA and trigger interferon response via cGAS/STING pathway(Simon *et al*., 2019). To measure the effects of S10 mutation on LINE1 expression we performed RNA sequencing on liver, brain, heart, and small intestine collected from WT, S10A and S10E mice. We then checked the expression of young and active LINE1 families(Sookdeo *et al*, 2013). We observed the trend of elevated LINE1 expression in small intestine in both male and female S10E mice and female S10E mice brain (**Fig. 2A, B**). Consistently with LINE1 inducing inflammation, in female brain, when comparing S10A and S10E mice to the WT mice, GSEA terms related to inflammation were downregulated in S10A mice and upregulated in S10E mice (**Fig EV. 1A**). However, in the female mice intestine which also showed upregulation of LINE1, the top enriched inflammation related GSEA terms showed the trend of upregulation in both S10A and S10E mice (**Fig EV.1B**). Interestingly, in the male mice intestine, the trend was opposite compared to the one in the female brain, the inflammation related enriched terms were upregulated in S10A mice but downregulated in S10E mice (**Fig EV. 1C**). As inflammation can be triggered by both elevated expression of LINE1 elements and increased DNA breaks the contribution of these two processes may differ in different tissues. We did not observe LINE1 expression changes in liver and heart probably due to the lower expression of SIRT6 in those organs (**Fig. 2A, B**).

**Figure EV1.**
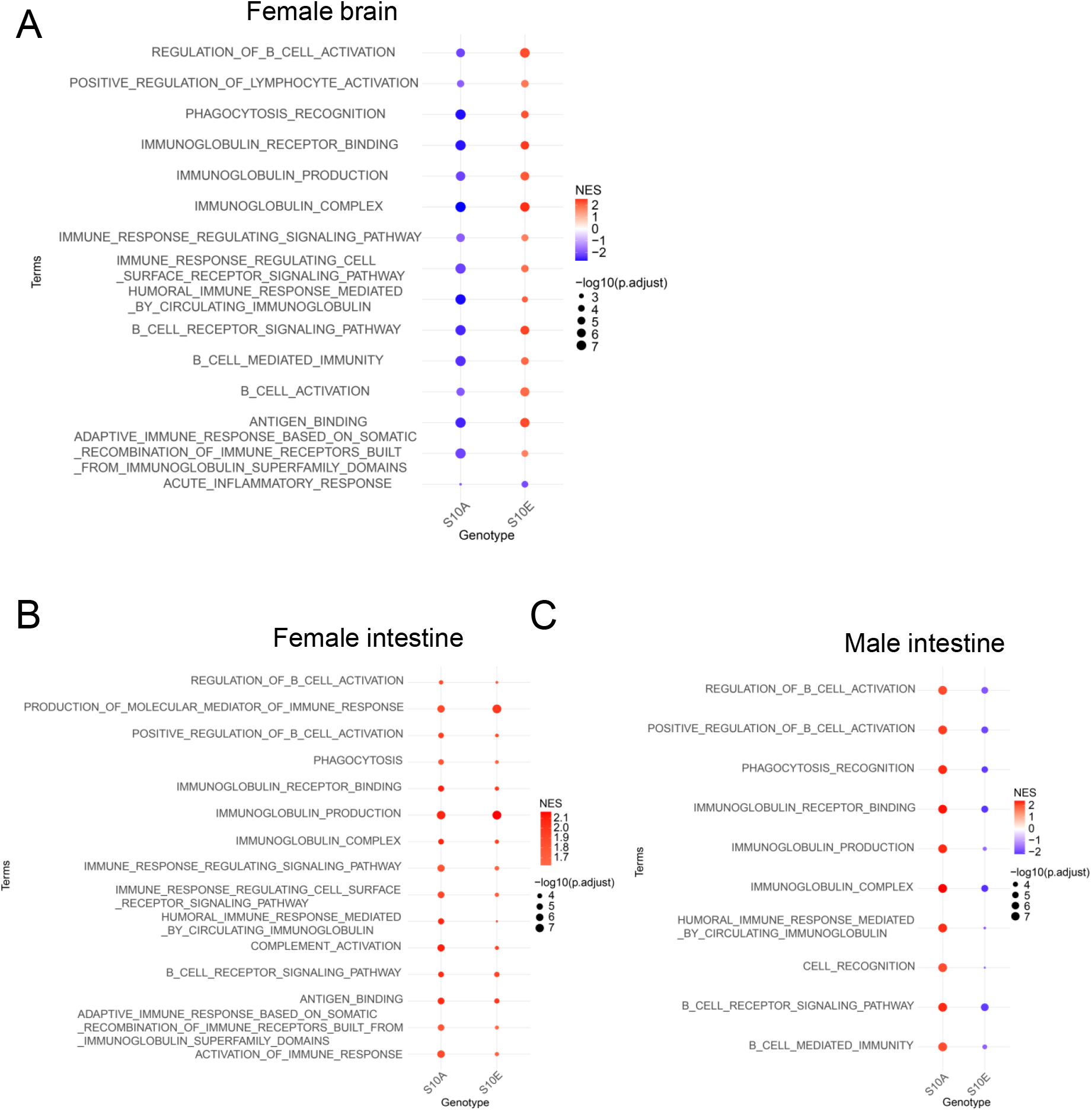
Immune and inflammation-related pathways are differentially altered by SIRT6 S10 mutations across tissues and sexes. Gene set enrichment analysis (GSEA) of immune and inflammation-related Gene Ontology biological process terms in RNA-seq data from young female (3-months-old) brains. Bubble plot shows normalized enrichment scores (NES; colour scale) and statistical significance (–log10(adjusted P); bubble size) for the top enriched terms when comparing S10A or S10E to WT mice. Terms include B-cell activation, immunoglobulin receptor signalling and acute inflammatory response. S10A brains generally show downregulation (negative NES, blue) of these pathways, whereas S10E brains show upregulation (positive NES, red), consistent with enhanced inflammatory signaling in the phosphomimetic mutants. Equivalent GSEA bubble plot for female small intestine. In contrast to the brain, most immune-related pathways are positively enriched in both S10A and S10E intestines relative to WT, suggesting that intestinal inflammation is increased in both mutants, with S10E showing the strongest effects. GSEA of immune-related pathways in male small intestine. Here, S10A intestines display positive enrichment of several B-cell and immunoglobulin-related pathways, whereas S10E intestines show negative or weaker enrichment, indicating that the balance between DNA damage–driven and LINE1-driven inflammatory signaling differs between sexes and tissues. In all panels, only pathways with adjusted P < 0.05 in at least one comparison are shown; NES and adjusted P values are derived from preranked GSEA using statistics.

**Figure 2.**
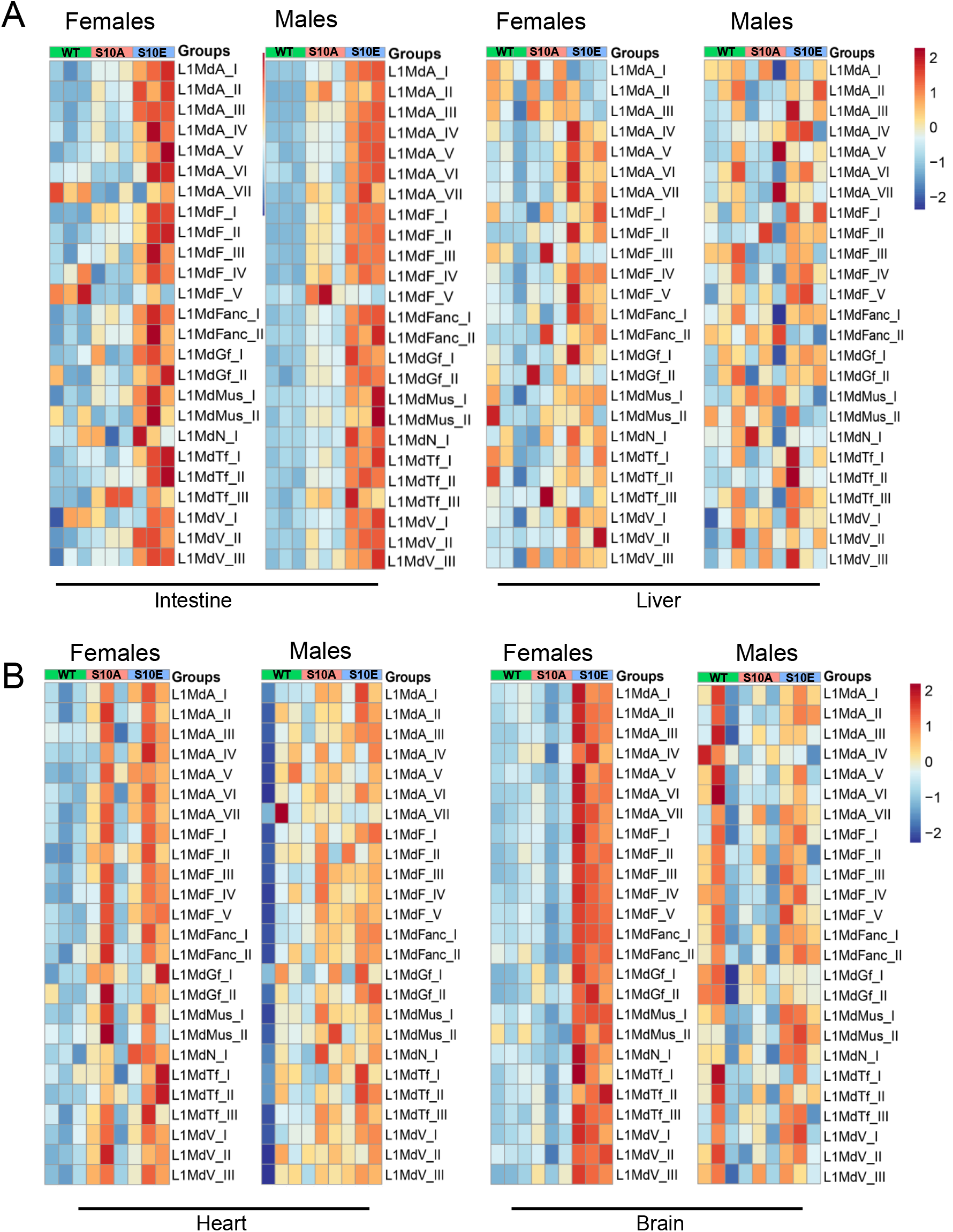
SIRT6 S10E mice exhibit tissue-specific upregulation of young LINE1 families. **A.** Heatmaps showing z-score–scaled RNA-seq expression of young and active LINE1 (L1Md) families in small intestine (left) and liver (right) from young (3-months-old) WT, S10A and S10E mice. Each column corresponds to an individual mouse and each row to a LINE1 family. Colors represent standardized expression (row-wise z-scores), with red indicating higher and blue indicating lower expression relative to the mean of that LINE1 family. Genotype groups (Young_WT, Young_S10A and Young_S10E) are indicated by color bars at the top of each heatmap. S10E mice show a consistent trend toward higher expression of multiple young LINE1 families in the intestine, whereas changes in liver are modest. **B.** Heatmaps of young LINE1 family expression in heart (left) and brain (right) from the same cohorts of mice. As in a, colors denote z-score–scaled expression. Female S10E brains display increased expression of several LINE1 families compared with WT, whereas changes in heart are comparatively subtle. Expression values were obtained from SalmonTE, and only LINE1 families previously annotated as young and potentially active were included.

### SIRT6 S10E mutant mice have shortened lifespan

The DNA damage response is widely recognized as a critical factor in aging and longevity(Burhans & Weinberger, 2007). Our previous research has also demonstrated that the capacity for DNA double-strand break repair co-evolves with maximum lifespan in rodents, with SIRT6 playing a key regulatory role in this process(Tian *et al*, 2019). Retrotransposons, such as LINE-1, are implicated as contributors to age-related pathologies and the aging process(De Cecco *et al*, 2019a; Simon *et al*., 2019). Given that SIRT6 is involved in both DNA repair and retrotransposon regulation, it is of particular interest to investigate how the two SIRT6 S10 mutants (S10A and S10E) impact the aging process in mice.

Interestingly, SIRT6 S10A mice of both sexes exhibited a lifespan comparable to the WT mice. However, S10E male were shorter-lived compared to both WT and S10A mice (**Fig. 3A**). S10E male mice exhibited a 10% and 14% median lifespan reduction compared to the WT and S10A mice, respectively. Both female and male S10E mice showed an increased body weight compared to the WT and S10A mice, which may be associated with increased inflammation or other metabolic changes (**Fig. 3B**).

**Figure 3.**
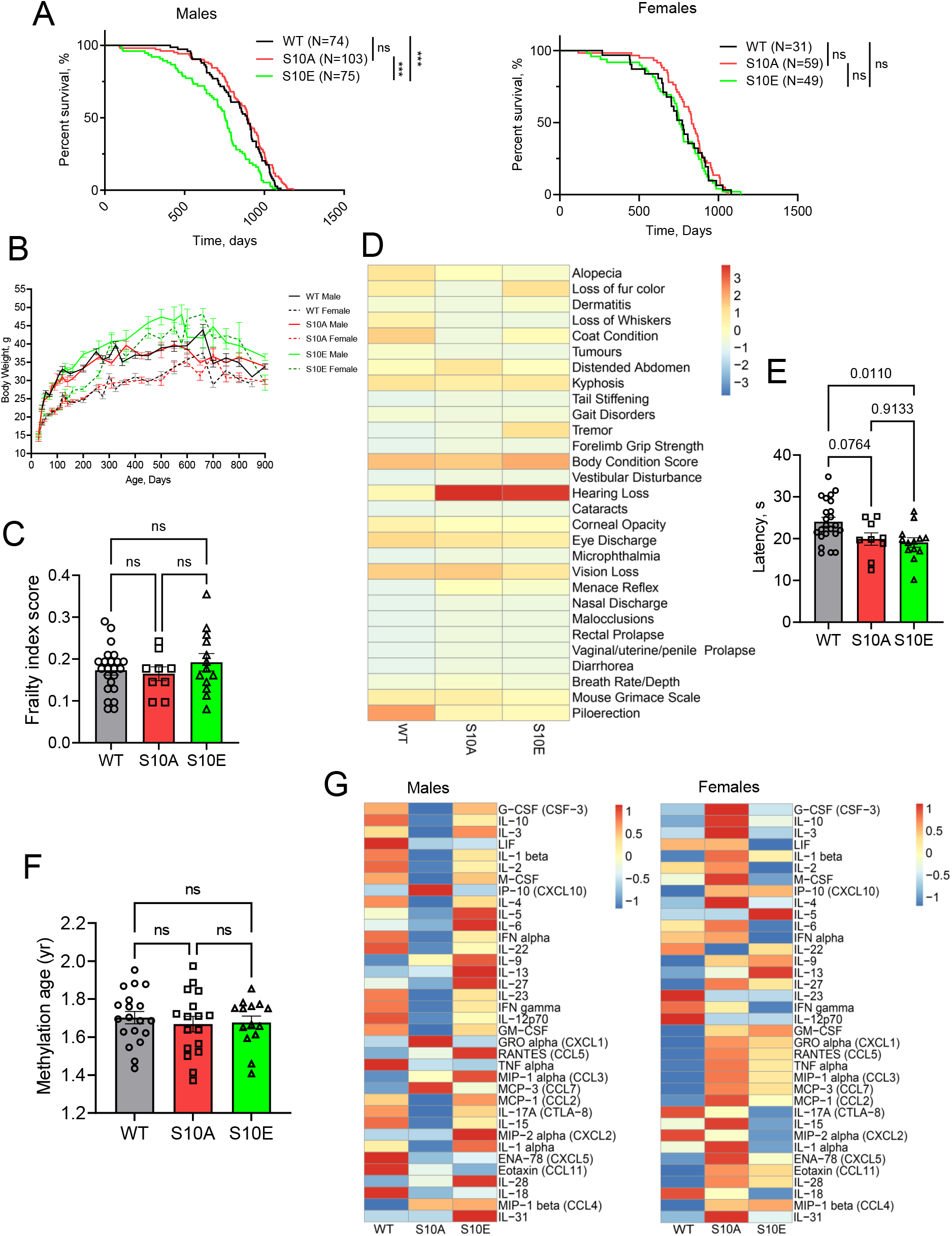
Lifespan and physiological phenotypes in SIRT6 S10A and S10E mice. **A.** Survival curves for WT, S10A and S10E mice separated by sex. Left, male mice (n shown on the plot); right, female mice (n shown on the plot). Mice were housed under identical conditions from weaning until natural death. S10E males show a significantly reduced median lifespan compared with WT and S10A males, whereas female lifespans are more similar across genotypes. Survival differences were analyzed using the Gehan– Breslow–Wilcoxon test. **\*\*\***: P<0.00. **B.** Longitudinal body-weight trajectories for WT, S10A and S10E mice of both sexes (n = X mice per genotype per sex). Points represent mean body weight at the indicated ages; error bars denote s.e.m. S10E mice of both sexes are heavier than WT and S10A mice across much of adult life, consistent with altered metabolic or inflammatory status. **C.** Frailty Index (FI) scores of aged mice (24 months old; n = 9-23 mice per genotype; sexes pooled). Each symbol represents one mouse; bars indicate mean ± s.e.m. The cumulative frailty burden is not significantly different among genotypes. P values were calculated by one-way ANOVA and are shown on the graph. **D**, Heatmap depicting the contribution of individual frailty parameters (for example, coat condition, kyphosis, tremor, grip strength, vision, vestibular function and hearing) to the overall FI in aged WT, S10A and S10E mice (n = 9-23 mice per genotype). Scores for each parameter are centered and scaled across animals (z-scores), highlighting specific deficits that tend to be more pronounced in S10A or S10E mice compared with WT. **E.** Rotarod performance of aged mice (n = 9-23 mice per genotype; sexes pooled). Latency to fall from an accelerating rotarod is shown for each mouse; bars represent mean ± s.e.m. Both mutant lines tend to perform worse than WT, with S10E mice showing the most pronounced deficit, indicative of impaired motor coordination and/or strength. P values were determined by one-way ANOVA with post-hoc tests and are indicated in the panel. **F.** Epigenetic age of liver DNA from 24-month-old WT, S10A and S10E mice, estimated using a mammalian DNA methylation clock (n = 14-19 mice per genotype). Each symbol represents one mouse; bars show mean ± s.e.m. No significant differences in methylation age are observed among genotypes at this time point. P values were calculated by one-way ANOVA. **G.** Heatmaps of plasma cytokine and chemokine levels measured by a 36-plex bead-based assay in 24-month-old WT, S10A and S10E mice (n = X mice per genotype per sex). Separate heatmaps are shown for males and females. Rows correspond to individual analytes (for example, IL-6, TNF, IFN-γ, MCP-1 and CXCL1), and columns to individual mice. Concentrations are centred and scaled per analyte (z-scores). Male S10A mice tend to show lower levels of several pro-inflammatory cytokines compared with WT and S10E males, whereas female S10A mice tend to show higher inflammatory cytokine levels.

To further evaluate the healthspan of these mice, we utilized a mouse frailty index(Whitehead *et al*, 2014), which integrates 31 parameters including body weight, temperature, coat condition, grip strength, mobility, vision, and hearing . Despite the comprehensive nature of this assessment, no significant differences were observed among the groups at 2 years of age (**Fig.3C**). However, upon closer examination of individual parameters, both aged S10A and S10E mice demonstrated a trend towards more severe hearing loss compared to the WT controls (**Fig. 3D**).

Locomotor function and coordination were evaluated using the rotarod performance test(Shiotsuki *et al*, 2010), where both S10A and S10E mice exhibited reduced time on the rotarod compared to Bl6 mice, though only the S10E group reached statistical significance (**Fig. 3E**).To assess biological age, we performed a methylation clock assay on DNA isolated from the livers of 2-year-old mice. Surprisingly, at this age, there were no significant differences in methylation age among the three genotypes (**Fig. 3F**).

To assess the inflammatory status of our mice, we performed a Luminex multiplex cytokine assay, which measures 36 different inflammatory cytokines and chemokines. Interestingly, male S10A mice exhibited reduced inflammation compared to the WT and S10E mice, while female S10A mice showed a trend toward increased inflammation. However, due to the substantial variability among individuals, none of the observed differences reached statistical significance (**Fig. 3G**). In summary, we show that S10E mutation shortens mouse lifespan in male mice, while S10A mutation shows a trend for longer median lifespan in female mice. As both DNA breaks and elevated LINE-1 expression result in increased inflammation, the balance between these processes may depend on the tissue and sex. Our results suggest that control of LINE1 expression has a stronger impact on mouse lifespan.

## Discussion

Serine 10 phosphorylation site on SIRT6 regulates SIRT6 recruitment to DSBs(Van Meter *et al*., 2016). It may also determine the balance between SIRT6 on DSBs versus LINE1 elements and other chromatin sites. Here we generated SIRT6 S10 mutant mice and analyzed how shifting this balance affects DNA repair and lifespan.

Consistent with the role of SIRT6 S10 phosphorylation in recruiting SIRT6 to DSBs (Van Meter *et al*., 2016), the S10E mice showed enhanced DNA repair capacity following γ-irradiation. Genomic instability is an important driver of ageing. DNA repair deficiency causes premature aging(Oshima *et al*, 2017; Rehman *et al*, 2020; Shiloh & Lederman, 2017) and long-lived species display more efficient DNA DSB repair(Firsanov *et al*, 2025; Tian *et al*., 2019).

However, the improvement in DNA repair in SIRT6 S10E mice did not translate into increased longevity. On the contrary, SIRT6 S10E male mice exhibited shortened lifespan compared to both WT and S10A mice. A possible explanation is that a benefit to lifespan in S10E mice was offset by deficiency in SIRT6 function in chromatin maintenance. Accordingly, in both male and female S10E mice, we observed increased LINE1 expression in several organs, consistent with previous findings that SIRT6 represses retrotransposon activity(Simon *et al*., 2019; Van Meter *et al*., 2014). Elevated LINE1 expression levels are known to trigger the cGAS/STING pathway, leading to sterile inflammation(De Cecco *et al*., 2019a; Simon *et al*., 2019; Van Meter *et al*., 2014) which can shorten lifespan.

There were no significant differences in frailty and methylation age between the three mouse genotypes, however, both S10A and S10E mice at 2 years of age displayed worse performance on rotarod and hearing loss indicating that impairment of either SIRT6 function affects mouse fitness.

While S10A mice showed reduced capacity to repair DNA breaks and reduced survival after irradiation, they displayed comparable lifespan to the WT mice in male mice, and a trend towards increased mean lifespan in female mice. This unexpected result suggests that SIRT6 function at repressing LINE1 elements, and possibly other chromatin related functions of SIRT6, are more important for mouse lifespan. While both DNA repair and LINE1 suppression are important for longevity, when the balance between these two SIRT6 functions is shifted, better chromatin maintenance outweighs enhanced DNA repair (**Fig. 4**). While our observations were limited to sedentary mice kept under standard vivarium conditions, it is possible that in the wild or under increased stress the relative impact of DNA repair and LINE1 suppression functions would be different.

**Figure 4.**
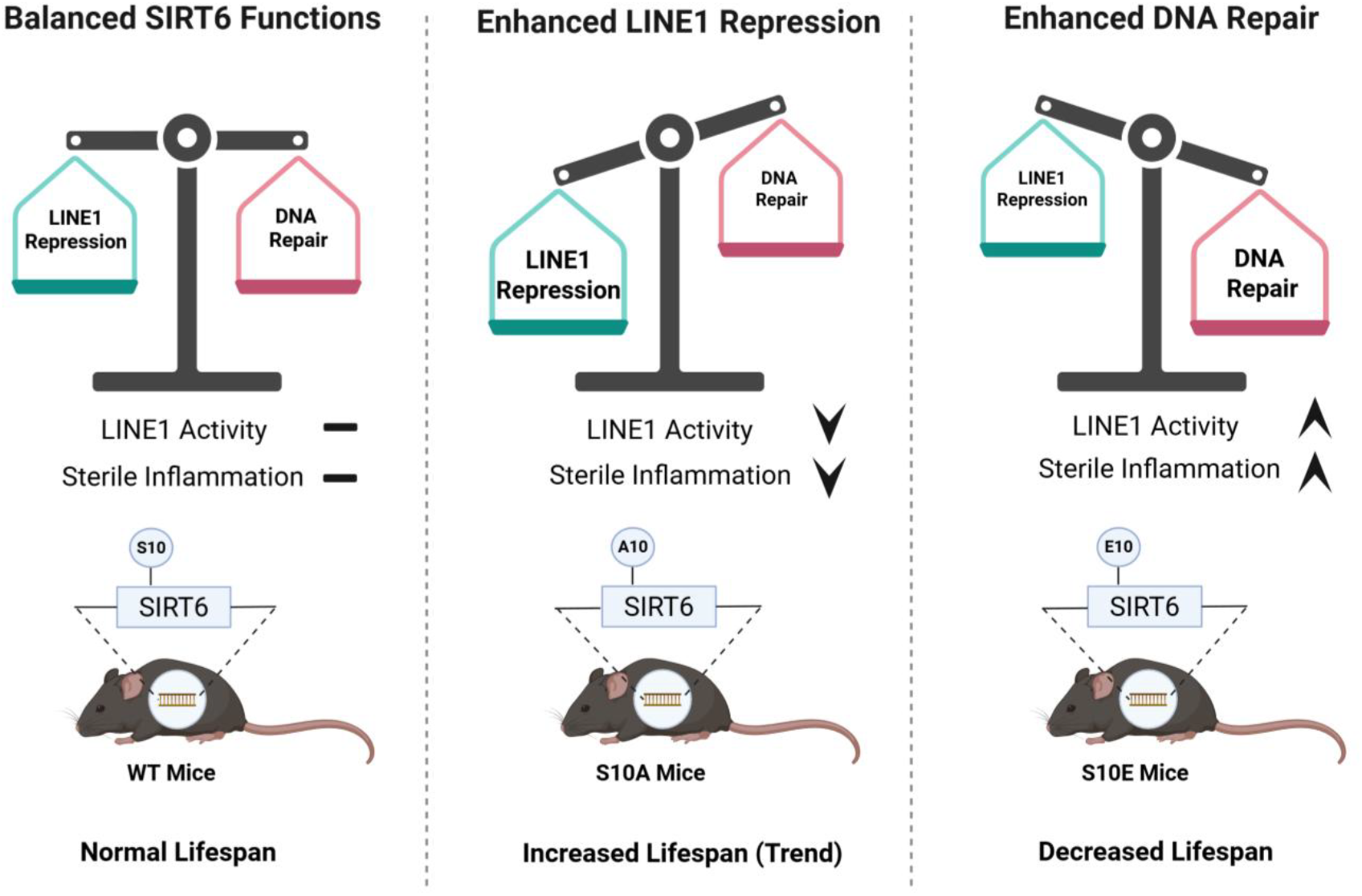
Schematic model illustrating how SIRT6 Ser10 status is proposed to tune the balance between LINE1 repression and DNA double-strand break repair, thereby modulating sterile inflammation and lifespan. In wild-type (WT) mice (left), SIRT6 bearing the native Ser10 (S10) maintains a functional equilibrium between LINE1 silencing and DNA repair, resulting in basal LINE1 activity, low sterile inflammation and a normal lifespan. In S10A mice (centre), the phospho-null A10 mutation is proposed to shift SIRT6 activity towards stronger LINE1 repression with relatively preserved DNA repair, leading to reduced LINE1 activity and decreased sterile inflammation, which may support increased lifespan. In S10E mice (right), the phosphomimetic E10 mutation is proposed to favour DNA repair functions at the expense of LINE1 repression; although DNA repair capacity is enhanced, loss of efficient LINE1 silencing results in elevated LINE1 activity and heightened sterile inflammation, which together are associated with a shortened lifespan.

In conclusion, our study highlights the multifaceted roles of SIRT6 Serine 10 phosphorylation in regulating DNA repair, LINE1, and inflammation. These findings suggest that targeting SIRT6 for therapeutic purposes will require a nuanced understanding of its tissue-specific and post-translational regulation.

## Acknowledgement

This work was supported by grants from the National Institutes of Health to V.N.G., A.S. and V.G (AG047200, AG092746, AG027237).

## Author contributions

Z.Z., M.S., A.S., and V.G. designed research, Z.Z., M.S., S.A.B., A.S., and V.G. analyzed data, and Z.Z., A.S., and V.G. wrote the manuscript; Z.Z., M.S. performed most of the experiments.

M.S. generated the transgenic mouse strain and performed with the aging study. Y.L. help with *in vivo* irradiation experiments. M.K. performed the immunofluorescence. V.P. helped with mouse colony maintenance and measured the frailty index score of mice. A.S. and V.G. supervised research.

## Methods

### Animal husbandry

All animal experiments were approved and performed in accordance with guidelines set forth by the University of Rochester Committee on Animal Resources with protocol number 2017-027 (mouse). Mice were group housed in IVC cages (up to 5 animals/cage) in a specific pathogen-free environment and fed standard chow diet (Altromin 1324; total pathogen free, irradiated with 25 kGy) and water ad libitum. Animal rooms were maintained at 21–24 °C and 35–75% relative humidity, with 12/12 h (6 a.m. to 6 p.m.) dark–light cycle. Cages were routinely replaced every 10–14 days.

### Generation of mutant mice

136nt ssODN containing either S10A or S10E mutations, along with silent PAM mutations, were designed and ordered through IDT. The sequences of the gRNA and two ssODN were listed below. Embryos were injected with ssODNs, along with Cas9 + Sirt6 sgRNA guides, and recovered into C57BL/6J surrogates. Formulation, injections, and generation of the transgenic mouse lines was facilitated by the Cornell University’s Stem Cell and Transgenic Core Facility. Founder animals were genotyped via sequencing upon receipt and backcrossed into stock C57BL/6J mice in reciprocal matings for 2 generations. Heterozygous animals were crossed in order to generate homozygous cohorts. Both S10A and S10E strains were maintained as homozygotes in colony. Both strains were backcrossed to C57BL/6J once per year before re-establishing homozygote breeders.

gRNA 5’-AGC AGG GTT GTC GCC TTA CG -3’

S10A ssODN:

5’-GGAAACTTTATTGTTCCCGTGCGGCAGCGCCGGCGACGATGTCGGTGAATTA

TGCAGCAGGGCTCGCTCCTTACGCAGATAAGGGCAAGTGCGGGCTGCCCGAGGT

GAGAGCTGCAGTGTTCGAGTCACCGAATGA -3’

S10E ssODN:

5’-GGAAACTTTATTGTTCCCGTGCGGCAGCGCCGGCGACGATGTCGGTGAATTA

TGCAGCAGGGCTCGAACCTTACGCAGATAAGGGCAAGTGCGGGCTGCCCGAGGT

GAGAGCTGCAGTGTTCGAGTCACCGAATGA -3’

### Whole body irradiation

Mice were exposed to a single sublethal dose of 9 Gy total body irradiation (TBI). Survival of mice was analyzed by Kaplan–Meier survival curves, and p-values were calculated by Gejam-Breslow-Wilcoxon test using Graphpad prism.

### Lifespan study

All the mice were housed in the same facility for all experiments. None of the animals entered into the aging study were allowed to breed. Mice were inspected daily for health issues, and any death was recorded. Animals showing significant signs of morbidity, based on the AAALAC guidelines, were euthanized for humane reasons and were used for lifespan analysis since they were deemed to live to their full lifespan. No mice were censored from analysis. Lifespan was analyzed by Kaplan–Meier survival curves, and p-values were calculated by Gejam-Breslow-Wilcoxon test using Graphpad prism.

### Tissue and plasma collection

Animals were brought to the laboratory in their holding cages and euthanized one by one for dissection. Mice were euthanized by isofluorane anesthesia followed by cervical dislocation. The dissection was performed as rapidly as possible following euthanasia by several trained staff members working in concert on one mouse. Tissue samples were either rapidly frozen in liquid nitrogen (for HA amount and MW determination, and RNA sequencing) or fixed in 4% formalin (for histology). Blood was collected by cardiac puncture into EDTA-coated tubes, centrifuged, and the plasma was aliquoted and rapidly frozen in liquid nitrogen. All frozen samples were stored at -80°C.

### Frailty index score

The Frailty Index (FI) was assessed as described previously(Whitehead *et al*., 2014). In brief, 31 health-related deficits were assessed for each mouse. A mouse was weighed, and body surface temperature was measured three times with an infrared thermometer (Fisher scientific). Bodyweight and temperature were scored based on their deviation from the mean weight and temperature of young mice(Whitehead *et al*., 2014). Twenty-nine other items across the integument, physical/musculoskeletal, ocular/nasal, digestive/urogenital, and respiratory systems were scored as 0, 0.5, and 1 based on the severity of the deficit. The total score across the items was divided by the number of items measured to give a frailty index score between 0 and 1.

### Rotarod performance

Motor performance was assessed using the protocol described before(Tung *et al*, 2014). Briefly, gross motor control was measured using the rotarod (IITC Life Science, CA, USA). For this test, each mouse was placed on a cylindrical dowel (69.5 mm in diameter) raised 27 cm above the floor of a landing platform. Mice were placed on the dowels for 5 min to allow them to acclimatize to the test apparatus. Once initiated the cylindrical dowels began rotating and accelerated from 5 rpm to a final speed of 20 rpm over 60 s. During this time, mice were required to walk in a forward direction on the rotating dowels for as long as possible. When the mice were no longer able to walk on the rotating dowels, they fell onto the landing platform below. This triggered the end of the trial for an animal and measurements of time to fall (TTF) were collected. Passive rotations where mice clung to and consequently rotated with the dowel were also used to define the end of the trial. Mice were then returned to their cages with access to food and water for 10 min. This procedure was repeated for a total of six trials, with the first three trials used for training and subsequent trials used for data analysis.

### RNA extraction for sequencing and qPCR

All frozen tissues were pulverized using the cell crusher. For preparing RNA from tissues, pulverized frozen tissues in the range of 10-15 mg were removed from samples kept at −80°C and extracted using Trizol reagent according to the supplier’s instructions. After recovery of total RNA from the Trizol reagent by isopropanol precipitation, RNA was digested with DNaseI for 30min at room temperature and further purified by PureLink™ RNA Mini Kit according to the instruction. The yield and quality were checked using Nano Drop. For PBMCs, RNA were purified using only PureLink™ RNA Mini Kit according to the instruction.

### RNA sequencing

The RNA samples were processed with the Illumina TruSeq stranded total RNA RiboZero Gold kit and then subjected to Illumina HiSeq 4000 paired-end 150bp sequencing at Univeristiy of Rochester Genomics Research Center. Over 50 million reads per sample were obtained. The RNA-seq experiment was performed in three biological replicates for all tissues. The RNA-seq reads were first processed using Trim_Galore (version 0.6.6), which trimmed both adapter sequences and low-quality base calls (Phred quality score < 20). The clean RNA-seq reads were used to quantify the gene expression with Salmon (version 1.4.0)(Patro *et al*, 2017). Specific parameters (--useVBOpt --seqBias --gcBias) were set for sequence-specific bias correction and fragment GC bias correction. Gencode(Frankish *et al*, 2019) (version M25) was used for the genome-wide annotation of the gene in the mouse. The reads counts for genes were used as the input for differential expression analysis by DESeq2(Anders & Huber, 2010). Low-expression genes with reads counts less than 10 were excluded. The cutoff for p-value and fold change was shown in the figure or figure legend. For LINE1 quantification, SalmonTE was run in quantification mode to generate both estimated counts and transcripts per million (TPM) values for each TE subfamily. LINE1 expression was then extracted by selecting all subfamilies annotated as LINE-1 (L1) and summing or analyzing them individually. For downstream analyses and visualization, we used the SalmonTE-reported counts for normalization and statistical testing and the TPM values (log_2_[TPM+1]) as a measure of relative LINE1 expression across samples and conditions.

Gene set enrichment analysis (GSEA)(Subramanian *et al*, 2005) was performed with “Preranked” model (version 4.1.0). All genes were preranked by the values of –log10 (adjusted p value)*(fold change)/abs(fold change). Adjusted p values and fold change were obtained from DEseq2. Normalization mode was set to “meandiv”. Only those gene set with a size more than 15 genes were kept for the further analysis.

### Cytokine assay

Thirty-six cytokines and chemokines were measured in 24 months-old mice plasma by luminex multiplex technique using a Cytokine & Chemokine 36-Plex Mouse ProcartaPlex™ Panel 1A kit (Thermo Fisher Scientific, cat no. EPX360-26092-901). The luminex multiplex assay was performed using undiluted plasma samples following the manufactures instruction.

### Methylation clock

Genomic DNA from 24 months old Bl6, S10A, and S10E mouse livers was purified using a DNeasy Blood & Tissue Kit (Qiagen, cat no. 69504). One hundred nanograms of purified genomic DNA were used for the methylation measurement. All DNAm data used was generated using the custom Illumina chip “HorvathMammalMethylChip40,(Arneson *et al*, 2021)” so-called the mammalian methylation array. The particular subset of species for each probe is provided in the chip manifest file can be found at Gene Expression Omnibus (GEO) at NCBI as platform GPL28271. The SeSaMe normalization method was used to define beta values for each probe(Zhou *et al*, 2018).

### qRT-PCR

For RT-qPCR, ∼300ng of purified RNA was revers transcribed into cDNA in the 20 μL using iScript cDNA synthesis kit. 2 μL of this reaction was used for subsequent qPCR reactions, which were performed using SYBR green system (Bio-Rad, cat. no. 1725124). The primer sequences were listed in table 1. The actin beta gene was used as internal normalization control.

**Table 1.**
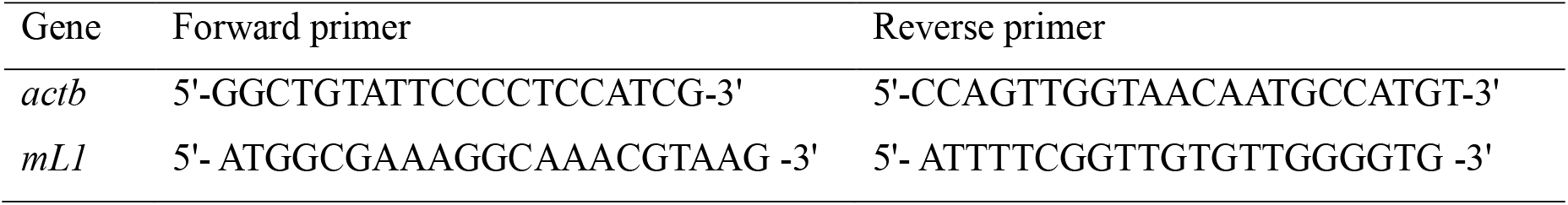

### γH2Ax staining of PBMCs

Mice were euthanized three days post-irradiation, and whole blood (150 µL) was collected via cardiac puncture for subsequent immunofluorescence staining. The collected blood was diluted at a 1:1 ratio with PBS containing 2% fetal bovine serum (FBS). Subsequently, 1.5 volumes of LymphoPrep solution were gently layered over the blood-PBS mixture. Peripheral blood mononuclear cells (PBMCs) were isolated by centrifugation at 800g for 20 minutes with the deceleration function disabled. The interphase containing the PBMCs was carefully collected, transferred to a new tube, and washed with PBS supplemented with 2% FBS. The washed cells were resuspended in 1 mL of PBS containing 2% FBS and attached to slides using a Rotofix 32A centrifuge. Cells were then fixed with 2% formaldehyde, permeabilized with 0.1% Triton X-100, blocked in 5% bovine serum albumin (BSA), and incubated overnight with an anti-phospho-Histone H2AX (Ser139) antibody (Sigma-Aldrich, 05-636, 1:1000). After washing with PBS-T, cells were incubated with Alexa Fluor 488-conjugated goat anti-mouse secondary antibody (Abcam, ab150113, 1:1000) for 1 hour at room temperature. Slides were mounted using VECTASHIELD Antifade Mounting Medium with DAPI and examined using a confocal microscope at ×100 magnification. Confocal images were acquired with a step size of 0.5 µm, spanning the full depth of the nuclei.

### Statistical and Demographic Analysis

Data are shown as means with SEM (unless stated otherwise). N indicates the number of animals per test group; age and sex are also noted. One-way annova test was used for all pairwise comparisons which satisfied with normal distribution. Mann-Whitney u test was used for data which is not satisfied with normal distribution. All relevant p values are shown in the figures. ns means no significance. Demographic data were processed with GraphPad Prism software to compute mean and median lifespans, SEM, percent increase of the median, and p values (log-rank test) for each cohort.

